# Prediction and Dissection of Nitrogenase Activity Based on Protein Sequences using Dual Descriptor

**DOI:** 10.64898/2026.11.18.767551

**Authors:** Yu-Jia Zhao, Gang Wu, Lin-Feng Wu, Xiao-Pan Hu, Feng-Biao Guo, Bin-Guang Ma

## Abstract

Nitrogenase reduces N_2_ to ammonia at ambient temperature and pressure, but its activity differs markedly among natural sequences, and conventional wet-laboratory screening suffers from low throughput and long turnaround times. Existing computational methods have partially alleviated this bottleneck, yet most of them rely on protein embeddings obtained from protein language models together with features such as gene expression, and their interpretability analyses mostly stop at the level of feature contribution. Here we present CarmnaDD, a purely sequence-driven framework built on the dual descriptor (DD): two learnable components, the composition weight map (CWM) and the position weight function (PWF), jointly generate a position-resolved description **N**(*k*), so that activity can be predicted without any prior knowledge beyond the sequence. On the same dataset used by Carmna, CarmnaDD attains *R*^2^ = 0.60 for regression and AUC = 0.94 and F1 = 0.86 for classification, better than our previous Carmna, which integrates multiple features. Interpretability analysis shows that the amplitude of the periodic terms of the PWF decreases as the rank increases, indicating that the positional encoding strategy adapts to the rank; the high-weight 3-mer fragments in the CWM are significantly enriched in polar uncharged residues, which corroborates the SHAP-based conclusion of Carmna; sliding-window scanning identifies fragments significantly associated with activity (*FDR* < 0.05), and the strongest window displays a D-X-E pattern with conserved flanks and a polymorphic center, sequences carrying R at the central position showing the highest activity; conservation is significantly negatively correlated (*ρ* = −0.30) with description intensity. Mapping the key windows onto the protein structure shows that four highly correlated windows converge on the nucleotide-binding interface and the conformational-transmission/protein-docking interface of NifH. These results provide a purely sequence-based framework for nitrogenase activity prediction that balances performance and interpretability.

## 1. Introduction

Ensuring global food security and promoting green and low-carbon agricultural transformation are core objectives of modern agricultural development. To continuously increase grain production, agricultural production has long relied heavily on synthetic nitrogen fertilizers. Although this has effectively alleviated food supply pressures, it has also brought about prominent resource depletion and ecological pollution issues [1]. Synthetic nitrogen fertilizer production is energy-intensive and associated with substantial upstream greenhouse gas emissions; moreover, field nitrogen use efficiency is less than 50%, with large amounts of unused nitrogen entering the environment, exacerbating climate change, soil acidification, and water eutrophication [2–4]. In contrast, biological nitrogen fixation mediated by nitrogenase, which converts atmospheric nitrogen into ammonia under ambient temperature and pressure, represents an ideal pathway for green agricultural production [5, 6].

Nitrogenase is the enzymatic basis of biological nitrogen fixation; its active site contains a structurally complex metallocofactor that cleaves the extremely stable N≡N triple bond under mild conditions [7]. Nitrogenases occur in three homologous forms, the molybdenum, vanadium and iron forms, among which molybdenum-based nitrogenase is the most widely distributed and the most thoroughly studied [8]. However, the complete picture of cofactor assembly and substrate reduction has not yet been elucidated [9–11]. Nitrogenase is extremely sensitive to oxygen, therefore, diazotrophs must maintain activity by strategies such as respiratory oxygen consumption, compartmentalization or conformational protection [12, 13]. From an applied perspective, natural nitrogenase sequences differ markedly in activity; however, conventional wet-laboratory screening suffers from low throughput, long experimental cycles, and high costs, making it difficult to support high-throughput mining of highly active nitrogen-fixing resources. Establishing an efficient computational approach for evaluating nitrogenase activity is therefore a key problem that urgently needs to be solved.

In recent years, machine learning has been driving protein science from trial and error towards data-driven discovery [14, 15]. At the level of enzyme function annotation, models such as DeepECtransformer exploit the Transformer architecture to achieve high-accuracy prediction of Enzyme Commission (EC) numbers [16]. At the level of enzyme kinetic parameter prediction, DLKcat takes protein sequences and substrate molecular graphs as input and directly predicts the enzyme turnover number *k_cat_* [17]; UniKP leverages pretrained protein language models to extend the prediction target to parameters such as *k_cat_*, *K_m_* and *k_cat_*/*K_m_* [18]; TurNuP characterizes the complete chemical reaction through differential reaction fingerprints and achieves cross-species *k_cat_* prediction for kinetically uncharacterized enzymes [19]; and CatPred integrates protein language model embeddings with three-dimensional structural information to achieve quantitative prediction of kinetic parameters such as *k_cat_*, *K_m_* and *K_i_* [20]. These advances show that deep learning can effectively capture the sequence–activity relationship of enzymes from sequence.

For the specific problem of nitrogenase activity prediction, Carmna provided the first systematic solution: it uses the pretrained protein language model ProtT5 to extract sequence embeddings [21] and integrates extra-sequence information such as dipeptide and pseudo-amino acid composition, gene expression levels, and codon preference to construct classification and regression models of nitrogenase activity, establishing the current performance benchmark for this task [22]. This approach nevertheless has limitations. First, the construction of extra-sequence features depends on corresponding expression profiles and genome annotations, which are often unavailable for newly sequenced strains with only genome sequences. Second, its interpretability analysis is limited to the ranking of feature importance.

To address these issues, we propose CarmnaDD, a framework for nitrogenase activity prediction based on the dual descriptor (DD) model [23]. The DD framework represents a protein sequence as the dual combination of two learnable parameter components: a composition weight map (CWM) that captures compositional features in units of *k*-mers, and a position weight function (PWF) that captures the position-dependent arrangement contributions through periodic basis functions. On this basis, CarmnaDD adopts a purely sequence-driven learning strategy to perform both regression and classification of nitrogenase activity. On the same dataset used by Carmna, CarmnaDD achieves *R*^2^ = 0.6035 for regression and AUC = 0.9377 and F1 = 0.8642 for classification, outperforming Carmna. Furthermore, we dissect the internal parameters of the CWM and the PWF, discuss how the model encodes positional information at different ranks, and relate the sequence fragments to which the model assigns high weights to known functional regions of nitrogenase (such as the P-loop and Switch I/II regions in NifH) [7, 24, 25], thereby linking predictive factors to biological mechanisms and providing a basis for interpreting nitrogenase activity.

## 2. Materials and methods

### 2.1 Data source, preprocessing and split

The nitrogenase dataset used in this study is exactly the same as that used by Carmna [22]. The dataset was compiled by Ye et al. and contains 402 nitrogenases, with activity measured by the acetylene reduction assay [26]. The data processing pipeline (**Figure 1A**) is briefly as follows: the amino acid sequences of the three subunits (nifD, nifH and nifK) are concatenated in the order nifD→nifH→nifK as the model input. Raw activity values (in nmol C_2_H_4_/mg protein/hour) are transformed by log_10_(activity + 2) before being used in the regression task; this transformation alleviates the skewed distribution of the data and avoids the numerical instability caused by zero or extremely small values. For the classification task, samples are divided into high-activity and low-activity classes using a threshold of 50 nmol C_2_H_4_/mg protein/hour. The training (321 sequences) and test (81 sequences) datasets are identical to that of Carmna. The training set is further divided into a training subset and a validation subset at a ratio of 4:1 randomly. For the classification task, stratified sampling is used to ensure balanced positive and negative samples in each subset.

**Figure 1.**
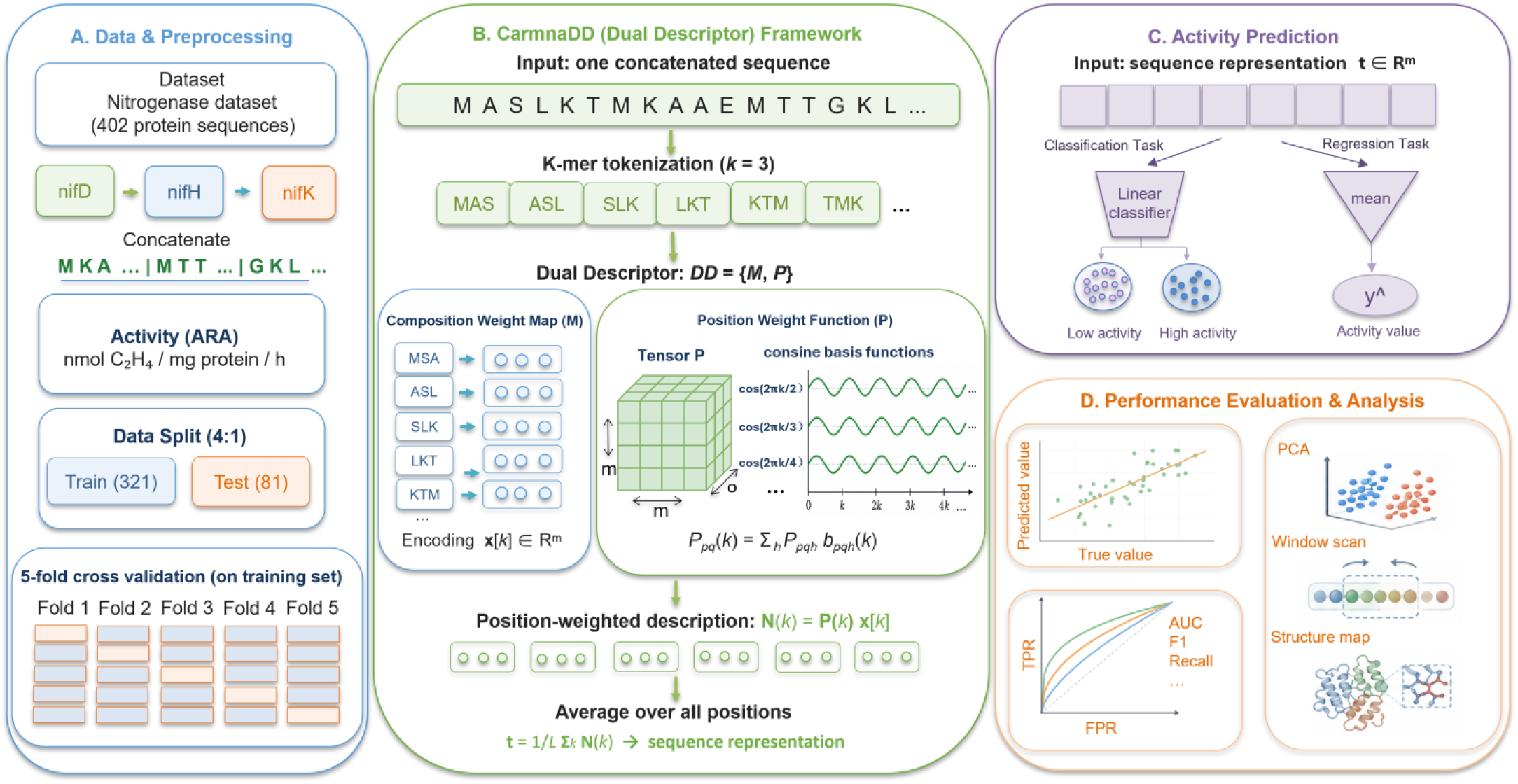
The workflow of CarmnaDD. (A) Data preprocessing and five-fold cross-validation. (B) Definition of Dual Descriptor (DD) for sequence representation. (C) Classification and Regression tasks for nitrogenase activity prediction. (D) Performance evaluation and interpretability analysis.

### 2.2 Dual descriptor (DD) model architecture

The core of CarmnaDD is the DD representation: *DD* = {*M*, *P*}, which decomposes sequence information into two complementary learnable components—the composition weight map (CWM, i.e., *M*) and the position weight function (PWF, i.e., *P*) [23]. The CWM describes the composition of the sequence tokens, and the PWF describes their position- or arrangement-dependent contribution (**Figure 1B**). Unlike conventional *k*-mer frequency representations, which encode only the occurrence frequency of each *k*-mer without explicitly modeling how the fragments are arranged, the DD representation characterizes both content and position.

Specifically, for an input protein sequence, the sequence is first divided into consecutive *k*-mer fragments (default *k* = 3 in this study). The DD rank *r* is defined as the fragment length, namely, the number of residues per token (*r* = 3 in this case), so that each token corresponds to one contiguous 3-mer and the vocabulary comprises 20^3^ possible tokens. The CWM *M* maps each fragment to an *m*-dimensional real-valued vector **x**[*k*], so that a sequence of length *L* is represented as the sequence of encoding vectors **x** = [**x**[1], **x**[2], …, **x**[*L*]].

The position weight function is implemented in tensor form: a learnable three-dimensional parameter tensor **P** ∈ ℝ*^m^*^×*m*×*o*^ is used, where *m* is the dimension of the encoding vector and *o* is the number of basis functions used by each position weight function. The basis functions are periodic cosine functions

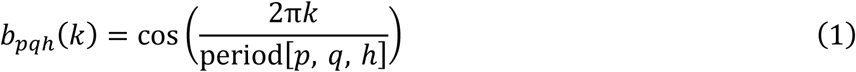

where *k* is the sequence position, *p* and *q* are the indexes in the encoding vector space (1 ≤ *p*, *q* ≤ *m*), respectively, and *h* is the basis function index (1 ≤ *h* ≤ *o*); the period is set by a fixed rule: **P**eriod[***P***, *q*, *h*] = ***P***(*m* · *o*) + *q* · *o* + *h* + 2. The (*p*, *q*)-th position weight function is 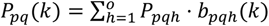. At position *k*, the position weight matrix **P**(*k*) is combined with the encoding vector **x**[*k*] to give the description at that position

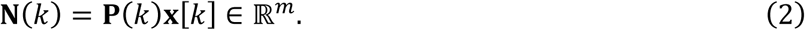

**N**(*k*) encodes both the compositional information of the *k*-mer at position *k* (from *M*) and its positional information in the sequence (from *P*), and is the basic unit of the DD framework. The sequence-level representation is the arithmetic mean over all positions **N**(*k*)

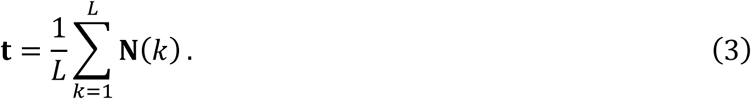

The regression task uses **t** to predict activity, with a sequence-level mean squared error loss; the classification task appends a linear classification head to **t** (**Figure 1C**). The specific hyperparameters of the classification and regression models are detailed in Sections 3.1 and 3.2.

### 2.3 Baseline models and evaluation metrics

To evaluate the performance of CarmnaDD, six classical machine learning algorithms were adopted as baselines, covering three classes of methods: kernel methods (support vector machine, SVM [27]; support vector regression, SVR [27]), neural networks (multilayer perceptron, MLP [28]) and tree ensembles (random forest, RF [29]; gradient boosting decision tree, GBDT [30]; light gradient boosting machine, LGBM [31]; extreme gradient boosting, XGBoost [32]). The baseline models SVM/SVR, MLP, RF, GBDT and all evaluation metrics were implemented with scikit-learn [33]; all baseline models used normalized 3-mer frequency vectors as input features. Classification models were evaluated by the area under the receiver operating characteristic curve (AUC), Precision, Recall and F1 score; regression models were evaluated by the coefficient of determination (*R*^2^), root mean squared error (RMSE), mean squared error (MSE) and mean absolute error (MAE). To ensure a fair comparison, the random seed for data splitting was fixed at 42 across all models (CarmnaDD and the baselines).

### 2.4 Model training and evaluation

Model training and evaluation followed the settings of Carmna, and the regression and classification tasks were trained independently. The Adam optimizer was used with an initial learning rate of 0.01, the learning rate was exponentially decayed by a factor of 0.99 per epoch, the batch size was 32 and the maximum number of epochs was 100; both tasks used five-fold cross-validation and early stopping, with the validation-set loss as the stopping criterion (training was terminated after 10 consecutive epochs without improvement, with a minimum improvement threshold of 1 × 10^−6^), and the model state with the lowest validation loss was retained; all performance evaluations were carried out on the independent test set.

### 2.5 Interpretability analysis

The interpretability analysis of CarmnaDD was carried out on the regression model (*rank* = 3, *vec_dim* = 26, *num_basis* = 18) (**Figure 1D**). All statistical tests were performed with a significance level of *α* = 0.05, and multiple comparisons were controlled for the false discovery rate (FDR) using the Benjamini–Hochberg procedure.

First, the two key components of the model were dissected. The L2 norm ||*M*|| of the embedding vectors of all 20^3^ = 8000 3-mers was calculated and sorted in descending order, and the top 20, 50 and 100 were taken as high-weight fragment sets; the residues contained in these fragments were extracted and counted according to five biochemical classes, with the amino acid composition of all residues in the training set (417,052 residues) used as background frequencies, and the enrichment fold was assessed by one-sided binomial tests with FDR correction. In parallel, models were trained under four rank settings (*rank* = 1–4; nonlinear mode), all elements of the parameter tensor **P** were extracted and sorted by absolute value, and the |*P*| distributions of the top 20 periodic terms were compared across the four settings.

Second, the DD representations at the sequence and subunit scales were analyzed. The arithmetic mean of **N**(*k*) over all positions of each sequence was taken as the sequence-level representation, on which principal component analysis (PCA) was performed; the principal components were then subjected to Pearson correlation analysis with nitrogenase activity. Similarly, PCA and correlation analysis were also performed on the sequence-level CWM representation (mean token embedding). Subunit intervals were defined in the order nifD → nifH → nifK using the annotated boundaries of each sequence; the mean description intensity of each subunit was computed as the arithmetic mean of **N**(*k*) over all its tokens; and differences among the three subunits were assessed with the Friedman test, followed by paired Wilcoxon signed-rank post-hoc comparisons, with each sequence treated as a block.

Finally, the residue-scale association with activity was examined. Multiple sequence alignment (MSA) of the 402 sequences was performed with the MUSCLE algorithm in MEGA11 (default parameters) [34, 35]. The description intensity at position *k* was defined as the L2 norm of **N**(*k*); windows of width 3 columns were slid along the alignment with a step of 1, and only strict windows without gaps in all 321 training sequences were retained (894 in total). The Pearson correlation coefficient between the local description intensity of each window and activity was computed, and significant windows were selected at *FDR* < 0.05 after Benjamini– Hochberg correction. In addition, the key windows were mapped onto the three-dimensional structure of nitrogenase (PDB: 1N2C, chain F [36]); the correspondence between MSA column numbers and structural residue numbers was determined from the sequence alignment; the minimum heavy-atom distance (excluding hydrogen atoms) from the residues of each window to the specified structural elements (P-loop, Switch I, Switch II, and 60’s loop) [24, 25], and to key ligands (ADP, [4Fe-4S] cluster, and Mg^2+^) [24, 25], as well as AlF_4_^-^ [36] was calculated. Subsequently, the Spearman rank correlation between the conservation of each strict window (defined as the mean frequency of the most common residue at the three sites) and its mean description intensity was computed, and description intensities were compared between conservation groups (≥ 0.9 and < 0.7) using the Mann–Whitney U test. The MSA file and the window scanning results are provided as **Supplementary Data** with this article.

## 3. Results and discussion

### 3.1 Classification model performance

To evaluate the performance of CarmnaDD on nitrogenase activity classification, we compared it with six classical machine learning algorithms under an identical data split. The hyperparameters of CarmnaDD for the classification task were set as follows: *rank* = 3, *vec_dim* = 30 and *num_basis* = 12. On the test set (**Table 1**), CarmnaDD achieved an AUC of 0.9377 and an F1 of 0.8642, with precision (0.8333) and recall (0.8974). Using the protein sequence as its only input, it outperformed Carmna (AUC = 0.9365, F1 = 0.8500), which additionally integrates multi-source information such as ProtT5 embedding features, gene expression profiles and codon preference. The ROC curves (**Figure 2**) further show that CarmnaDD has the largest area under the curve and maintains a higher true positive rate across all threshold ranges. This indicates that the DD model can extract discriminative features related to enzyme activity directly from the raw amino acid sequence. Among the six baseline models, the multilayer perceptron (MLP) achieved the highest AUC (0.9072) and F1 (0.8605), but its precision (0.7872) and recall (0.9487) were clearly imbalanced, indicating that the model tends to predict more samples as positive; the tree-based ensemble algorithms (XGBoost, GBDT, LGBM and RF) performed worse overall than MLP and SVM, possibly because the 3-mer frequency feature space is high-dimensional and sparse.

**Figure 2.**
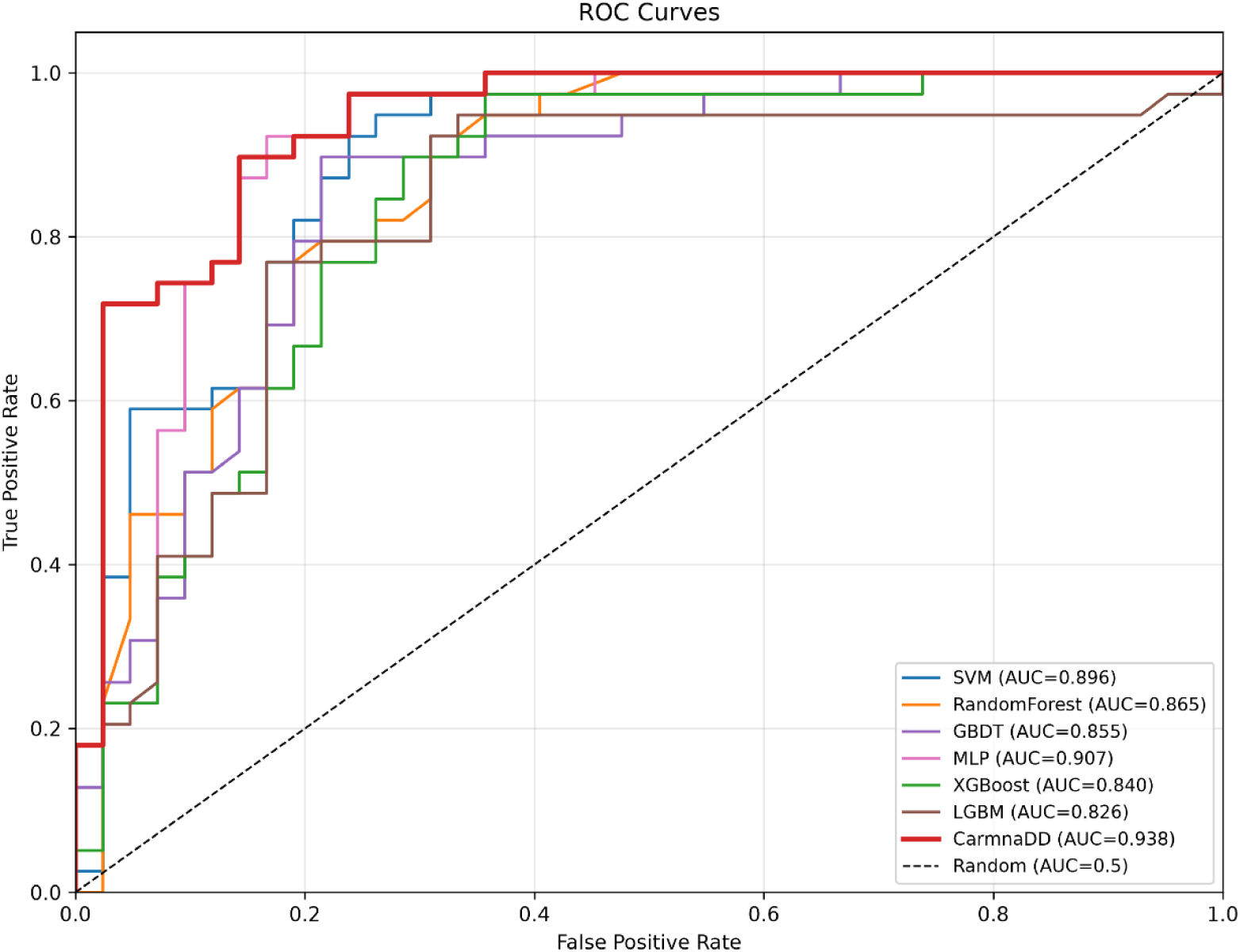
ROC curves of various models for nitrogenase activity classification. The *x*-axis is false positive rate and the *y*-axis is true positive rate. The AUC of each model is shown in the legend, and the black dashed line represents the random prediction baseline (AUC = 0.5).

**Table 1.** Classification task performance on the test set.

| **Model** | **AUC** | **Precision** | **Recall** | **F1** |
|---|---|---|---|---|
| SVM | 0.8962 | 0.7826 | 0.9231 | 0.8471 |
| MLP | 0.9072 | 0.7872 | **0.9487** | 0.8605 |
| RF | 0.8651 | 0.7895 | 0.7692 | 0.7792 |
| GBDT | 0.8550 | 0.7949 | 0.7949 | 0.7949 |
| LGBM | 0.8260 | 0.7895 | 0.7692 | 0.7792 |
| XGBoost | 0.8400 | 0.7568 | 0.7179 | 0.7368 |
| Carmna | 0.9365 | 0.8293 | 0.8718 | 0.8500 |
| CarmnaDD | **0.9377** | **0.8333** | 0.8974 | **0.8642** |

### 3.2 Regression model performance

To evaluate the quantitative prediction performance of CarmnaDD for nitrogenase activity, we first systematically validated the regression performance of the model. The hyperparameters of CarmnaDD for the regression task were uniformly set as *rank* = 3, *vec_dim* = 26 and *num_basis* = 18. Under standard five-fold cross-validation and a unified data split, CarmnaDD achieved excellent prediction on the test set, with a determination coefficient *R*^2^ of 0.6035, substantially better than the control models and the earlier algorithm. Compared with the best conventional baseline model, SVR, CarmnaDD improved *R*^2^ by 0.063; compared with the earlier Carmna model (*R*^2^ = 0.5572), it improved by 0.046. Its RMSE was 0.5381 and its MAE 0.3205, the lowest error indices among all compared methods, demonstrating its advantage in accurate and stable quantitative prediction. The scatter plots of predicted versus true values (**Figure 3**) further confirm that the predictions of CarmnaDD closely follow the diagonal of perfect fit and maintain good linear agreement even in the medium-to-high activity range of nitrogenase, indicating that the DD framework can mine and extract more comprehensive and more relevant enzyme activity features from raw protein sequences and effectively overcomes the representational limitations of conventional sequence features. Six mainstream regression models were also used as baselines, covering kernel methods (SVR), ensemble learning algorithms (RF, GBDT, XGBoost and LGBM) and a neural network model (MLP), all of which used the unified 3-mer frequencies as input features. The results (**Table 2**) show that the test-set *R*^2^ of the six baseline models ranged from 0.4625 to 0.5405, with SVR performing best, followed by XGBoost and GBDT, while MLP, RF and LGBM performed relatively poorly. This result indicates that, although conventional 3-mer frequency features can capture the compositional information of protein sequences to a preliminary extent, they suffer from pronounced high-dimensional sparsity, and relying on a single type of compositional feature cannot fully or accurately represent nitrogenase activity, which greatly limits the quantitative prediction ability of conventional machine learning models and further confirms the superiority and effectiveness of the feature extraction and modeling strategy of CarmnaDD.

**Figure 3.**
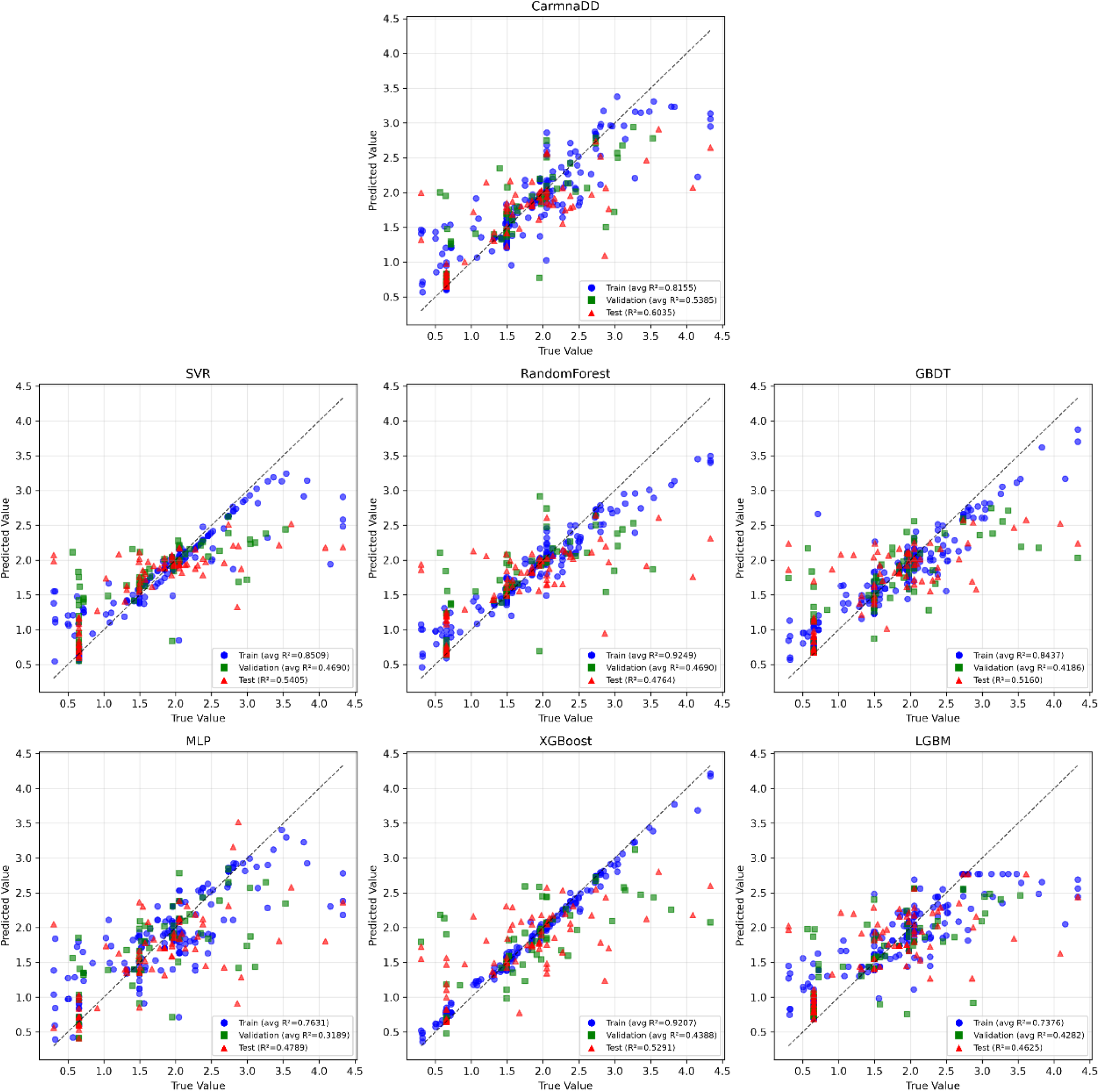
Scatter plots of predicted versus true nitrogenase activity by different regression models. Blue circles, green squares and red triangles correspond to training, validation and test samples. The diagonal dashed line indicates perfect prediction, and average *R*^2^ of each dataset is labeled in each subplot.

**Table 2.** Regression task performance on the test set.

| **Model** | **R<sup>2</sup>** | **RMSE** | **MSE** | **MAE** |
|---|---|---|---|---|
| SVR | 0.5405 | 0.5793 | 0.3356 | 0.3616 |
| MLP | 0.4789 | 0.6169 | 0.3805 | 0.3686 |
| RF | 0.4764 | 0.6184 | 0.3824 | 0.3782 |
| GBDT | 0.5160 | 0.5946 | 0.3535 | 0.3893 |
| LGBM | 0.4625 | 0.6265 | 0.3925 | 0.4096 |
| XGBoost | 0.5291 | 0.5864 | 0.3439 | 0.3794 |
| Carmna | 0.5572 | 0.5687 | 0.3234 | 0.3351 |
| CarmnaDD | **0.6035** | **0.5381** | **0.2896** | **0.3205** |

### 3.3 Interpretability analysis of the model parameters

#### 3.3.1 Interpretation of the CWM

In the CarmnaDD model, the encoding matrix **M** holds the learnable parameters of the composition weight map (CWM), which assigns a vector representation of the same dimension to each *k*-mer token. For the regression setting with *rank* = 3, there are 20^3^ = 8000 amino acid 3-mer fragments, each corresponding to a learnable embedding vector. To assess the relative contribution of different 3-mer fragments to the model, the L2 norm of each embedding vector was calculated and all 3-mers were ranked by norm. Under the nonlinear mode with *rank* = 3, the top 20 3-mers with the highest ||*M*|| were obtained (**Figure 4A**). The CWM parameters **M** mainly capture amino acid composition patterns that contribute strongly to the sequence representation; 3-mer fragments with high norms such as ASN, EHN, EKP and YIK possess larger embedding norms, indicating that these short peptide fragments have a stronger influence on the sequence-level representation learned by the model.

**Figure 4.**
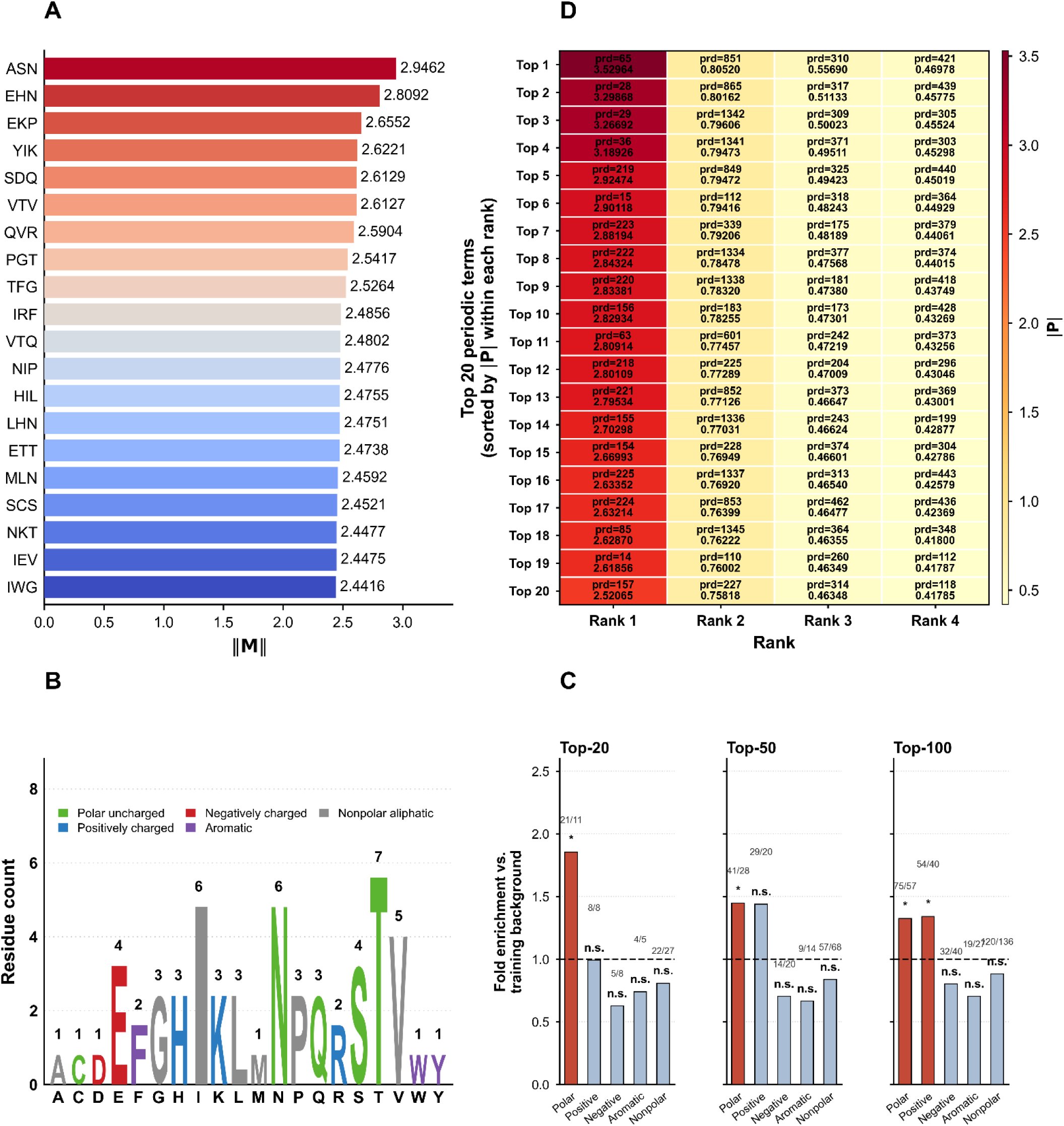
Analysis of high-weight 3-mer tokens and periodic terms in CarmnaDD. (A) Horizontal bar plot showing the top-20 amino-acid 3-mers ranked by the L2 norm ||M|| of CWM embedding vectors under nonlinear mode with *rank* = 3. Bar colors are rendered from blue to red according to the rank of ||M||, where red indicates a higher rank. (B) Residue-letter bar plot for the Top-20 high-weight 3-mer set. Residues are classified into five biochemical groups: polar uncharged (S, T, N, Q, C), positively charged (K, R, H), negatively charged (D, E), aromatic (F, W, Y), and nonpolar aliphatic (A, G, I, L, M, P, V). Each residue group is indicated by a distinct color. (C) Amino acid enrichment in top CWM 3-mers. Fold enrichment of the five biochemical residue classes in the Top-20 (60 residues), Top-50 (150 residues), and Top-100 (300 residues) highest-norm 3-mer sets, relative to the background amino acid composition of the 321 training sequences (417,052 residues). Red bars indicate significant enrichment (one-sided binomial test, Benjamini–Hochberg FDR *q* < 0.05); n.s., not significant. Numerals above the bars give the observed/expected residue counts. (D) Distribution of |P| values for periodic terms in the PWF of CarmnaDD across different rank settings (nonlinear mode). The heatmap shows the top 20 periodic terms sorted by the absolute value of the elements in tensor **P** (|P|) across ranks 1 to 4. Each cell displays the corresponding period index (prd) and its |P| value.

To dissect the residue composition preference within the high-norm 3-mers, all amino acid residues contained in the top-ranked 3-mers were extracted and analyzed statistically according to five biochemical classes (polar uncharged, positively charged, negatively charged, aromatic, nonpolar aliphatic). Taking the amino acid composition calculated from the 417,052 residues of the 321 training sequences as the background, the background frequencies of the five classes were 18.89%, 13.44%, 13.30%, 9.01% and 45.36%, espectively. The residue distribution of the top-20 high-weight 3-mer set is visualized by a residue-letter bar plot (**Figure 4B**). One-sided binomial tests were used to assess the residue enrichment folds, and the Benjamini–Hochberg procedure was used for multiple-testing correction to control the FDR; the enrichment statistics are given in **Figure 4C**. The statistical results show that polar uncharged residues were significantly enriched in all three high-weight fragment sets (**Figure 4C**): top-20 (60 residues, 21/60, enrichment fold 1.85, *q* = 0.0119), top-50 (150 residues, 41/150, enrichment fold 1.45, *q* = 0.0365) and top-100 (300 residues, 75/300, enrichment fold 1.32, *q* = 0.0266). This enrichment of polar uncharged residues corroborates the conclusion of Carmna based on SHAP analysis, which proposed that increasing the content of five amino acids, namely Ala, Asn, Gln, Thr and Val, promotes nitrogenase activity [22]. At the same time, an enrichment trend of positively charged residues (K, R and H) was observed, which is consistent with the understanding of nitrogenase structural biology: positively charged residues (such as lysine, arginine, and histidine) can mediate electrostatic interactions that participate in FeMo-cofactor assembly [37] and the mutual recognition of the Fe protein and the MoFe protein [38]; moreover, specific basic residues (e.g., arginines) along the water chain contribute to the electrostatic/hydrogen-bonding microenvironment essential for proton–electron transfer within the catalytic pocket [39]. It should be noted that although a positive enrichment trend was observed across the top-50 and top-100 sets, the enrichment of positively charged residues was statistically significant only in the top-100 set after multiple-testing correction. Therefore, its statistical robustness is limited, and the functional significance suggested here still requires further validation by functional experiments such as site-directed mutagenesis.

#### 3.3.2 Interpretation of the PWF

The PWF encodes positional information into the sequence representation through the learnable parameter tensor **P** and periodic cosine basis functions. To examine the differences in how the model exploits positional information under different rank settings, the |*P*| distributions of the top 20 periodic terms were compared for *rank* = 1–4 (nonlinear mode, non-overlapping token extraction) (**Figure 4D**). The results show two main trends: first, the overall magnitude of |***P***| decreases as the rank increases, and at *rank* = 1 the |***P***| of the top 20 terms is markedly larger than at the other ranks; second, the relative ranking of the top periodic terms is rearranged among ranks. These observations suggest that the PWF adopts differentiated encoding strategies at different granularities: at *rank* = 1, each token covers only a single residue and the available local sequence information is limited, so the model must rely on periodic terms of larger amplitude to enhance the expressiveness of the positional encoding; as the rank increases, the tokens themselves carry richer sequence information, and the dependence on periodic enhancement decreases accordingly.

### 3.4 Interpretability analysis of description granularity

#### 3.4.1 Interpretation of the sequence-level representation

Averaging **N**(*k*) over all positions of each sequence gave a 26-dimensional sequence-level representation, on which PCA was performed (**Figure 5A**). The first principal component (PC1) explained 98.5% of the variance and was highly correlated with activity ( *r* = +0.8865, ***P*** = 9.3 × 10^−109^); high-activity (upper tertile, *n* = 118) and low-activity (lower tertile, *n* = 146) sequences were significantly separated along PC1 (Mann–Whitney U test, ***P*** = 9.3 × 10^−42^). The second to fourth principal components contributed only 0.8%, 0.4% and 0.2% of the variance, respectively, and none of them was significantly correlated with activity (***P*** ≥ 0.13). These results indicate that the **N**(*k*) representation concentrates activity information along a single direction, forming an “activity axis”.

**Figure 5.**
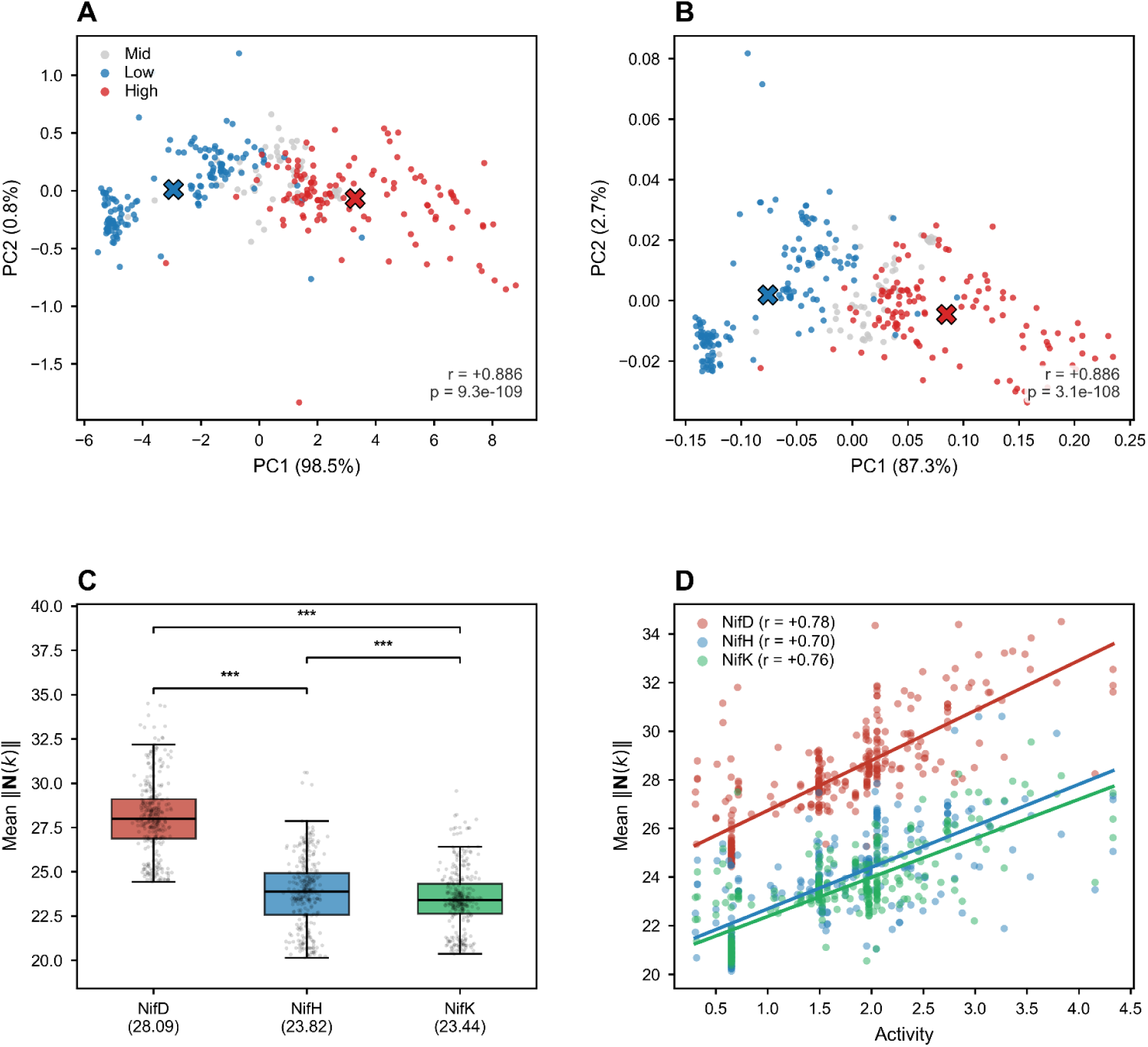
Principal component structure of the sequence-level representations and allocation of subunit description intensity. (A) PCA of the sequence-level **N**(k) representation. PC1 accounts for 98.5% of the variance and correlates strongly with activity (*r* = +0.8865, *p* = 9.3×10^-109^). Red, grey and blue denote high (*n* = 118), medium (*n* = 57) and low (*n* = 146) activity groups; ✕ marks the group centroids of the high and low groups. (B) PCA of the sequence-level CWM representation. PC1 accounts for 87.3% of the variance (*r* = +0.8855, *p* = 3.1×10^-108^); PC2 (2.7%) is unrelated to activity (*r* = +0.071, *p* = 0.207) but correlates with residue composition. Grouping as in (A). (C) Mean subunit description intensity (*n* = 321). Box plots show median and interquartile range; grey dots represent individual sequences. NifD 28.089 ± 2.154 > NifH 23.816 ± 2.001 > NifK 23.437 ± 1.739; all pairwise comparisons are significant (*p* < 10^-3^). (D) Description intensity of each subunit versus activity; lines are linear fits.

The same analysis was performed on the sequence-level CWM representation (mean token embedding) (**Figure 5B**). PC1 explained 87.3% of the variance and correlated with activity almost as strongly as the **N**(*k*) representation ( *r* = +0.8855, ***P*** = 3.1 × 10^−108^). The second principal component (PC2, 2.7%) was unrelated to activity (*r* = +0.071, ***P*** = 0.207) but was significantly correlated with amino acid composition: it was positively correlated with the proportion of positively charged residues (K, R and H) (*r* = +0.671, ***P*** = 2.1 × 10^−43^). These results indicate that, while encoding activity information, the CWM representation independently carries amino acid composition information in its low-variance directions. The positive-charge correlation revealed by PC2 corroborates the enrichment analysis based on token embedding norms, which likewise found that the positive-charge residues were enriched in the high-weight 3-mer sets (**Figure 4C**). The two analyses start independently from the variance structure at the sequence level and from the weight ranking at the token level, and point to a consistent conclusion.

#### 3.4.2 Allocation of subunit description intensity

The mean description intensity of each subunit was calculated in the coding order nifD → nifH → nifK (*n* = 321; **Figure 5C**). The mean description intensities of the three subunits were, in order, NifD 28.089 ± 2.154, NifH 23.816 ± 2.001 and NifK 23.437 ± 1.739; the overall difference was highly significant and all pairwise comparisons were also significant (Friedman *χ*^2^ = 484.5, *df* = 2, *p* = 6.2 × 10^-106^; paired Wilcoxon, all pairwise *p* < 1×10^-6^). The description intensities of the three subunits were all significantly positively correlated with activity (NifD *r* = 0.782, NifH *r* = 0.699, NifK *r* = 0.757; all *p* < 1×10^-47^; **Figure 5D**). One possible explanation is that the allocation of description intensity is determined solely by sequence length. However, the measured subunit lengths were NifK (514.9 ± 9.8 aa) > NifD (490.0 ± 10.4 aa) > NifH (294.6 ± 16.1 aa), which is inconsistent with the order of intensities (NifD > NifH > NifK)—the longest subunit, NifK, instead had the lowest intensity. These observations argue against sequence length as the driver of intensity allocation. Structural biology studies of nitrogenase have clearly established the division of function among the three subunits: the MoFe protein is an α_2_β_2_ heterotetramer whose α subunit (NifD) contains the FeMo-cofactor and is the catalytic center for substrate binding and reduction [40]; the β subunit (NifK) together with the α subunit coordinates the P-cluster at their interface [41]; and the Fe protein (NifH) is a homodimer containing a bridging [4Fe–4S] cluster and a pair of MgATP-binding sites [24]; it is responsible for transferring electrons to the MoFe protein, and the dissociation of its complex with the MoFe protein is the rate-limiting step of the entire catalytic cycle [42]. We observed that, without using any structural information, the model assigned the highest mean description intensity to the NifD subunit, which harbors the actual catalytic site, consistent with the functional importance of this subunit [40].

#### 3.4.3 Identification of activity-associated sites

To identify local sites in the sequence that are associated with nitrogenase activity, a sliding-window scan with a window of 3 columns was performed on the multiple sequence alignment (321 training sequences), yielding 894 strict windows that satisfied the criterion of no gaps within the window. For each window, ‖**N**(*k*)‖ of the 3-mer token corresponding to that window was taken as the local description intensity of the window, its Pearson correlation coefficient with activity was calculated, and FDR correction was then applied with the Benjamini–Hochberg method. A total of 484 windows (54.1%; **Figure 6A**) were significant at *FDR* < 0.05, and the top-ranked windows were concentrated in the 950–1050 region of the MSA (such as the strongly positively correlated windows 977 and 994, and the strongest negatively correlated window 1026), suggesting that this segment carries the most concentrated activity-related signal.

**Figure 6.**
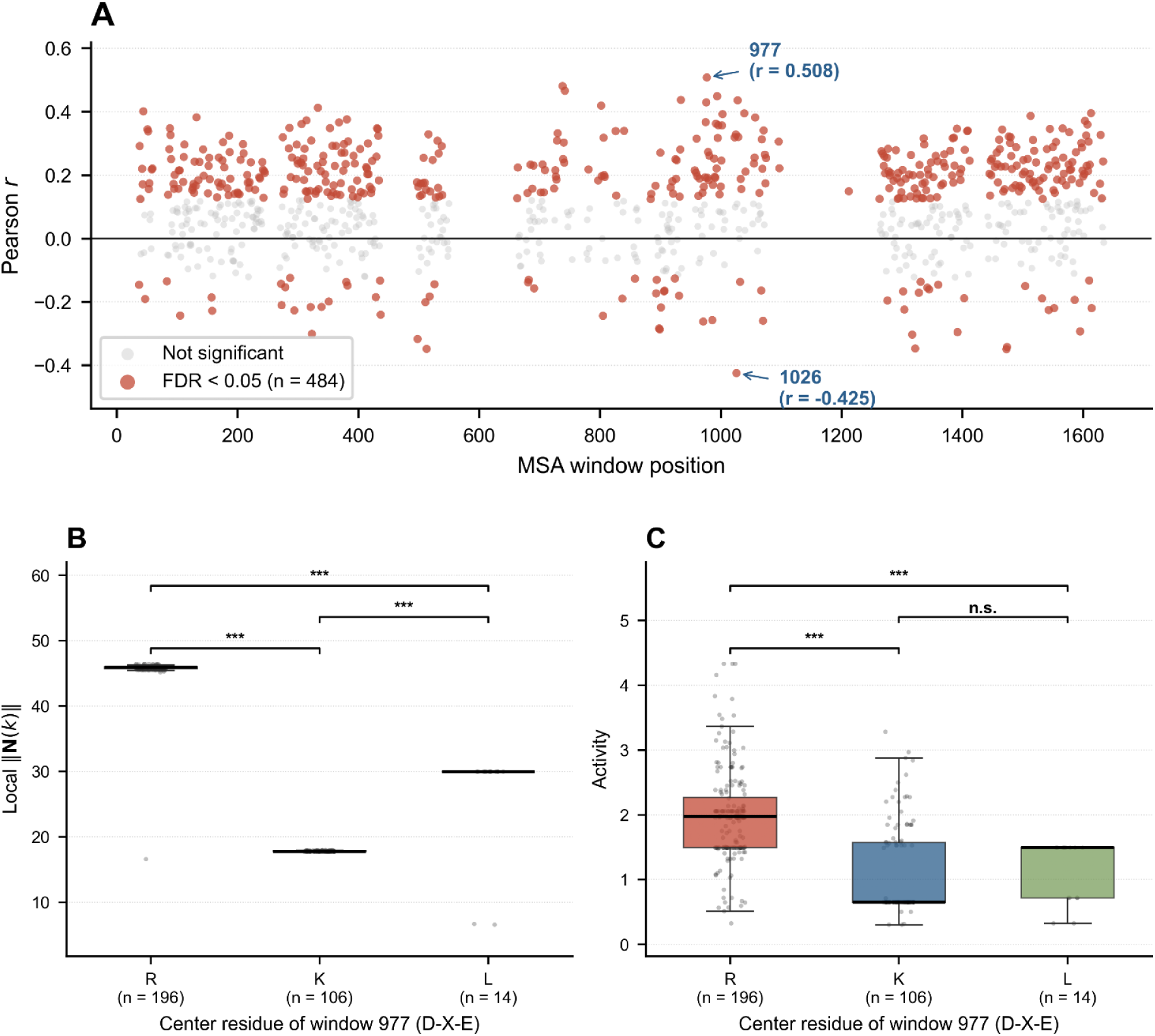
Identification of activity-associated sites. (A) Distribution of correlation coefficients from the MSA window scan. The *x*-axis denotes the window position as the column index in the MSA (0-based); the *y*-axis denotes the Pearson correlation coefficient *r* between the local description intensity of the window and nitrogenase activity. Significant windows are shown as filled markers (*FDR* < 0.05, Benjamini–Hochberg correction) and non-significant windows in grey. (B) Distribution of the local description intensity ‖**N**(*k*)‖ for the center residue (R, K, L) at window 977. (C) Distribution of nitrogenase activity for the same three groups.

Window 977 shows a typical “conserved flanks, variable center” pattern: the aspartate (D) at the first position is present in 318/321 sequences (conservation 0.991) and the glutamate (E) at the third position is present in all sequences (conservation 1.000), whereas the central second position is polymorphic, mainly R (*n* = 196), K (*n* = 106) and L (*n* = 14), with a few other types such as G (*n* = 4) and E (*n* = 1). After grouping by the central residue, the local description intensity differed highly significantly among the three groups (Kruskal–Wallis, ***P*** < 10^−16^), and activity also differed significantly ( ***P*** = 1.5^6^ × 10^−18^). Notably, the ordering of the local description intensity and of activity across groups was not consistent: the intensity order was R > L > K, whereas the activity order was R > K > L (**Figure 6B, C**). Pairwise comparisons further confirm this—R versus K and R versus L differed significantly in both intensity and activity, whereas K versus L differed significantly in intensity (***P*** = 2.77 × 10^−6^) but not in activity (***P*** = 0.441). This indicates that, although the local description intensity is overall correlated with activity, it is not sufficient to characterize activity on its own.

From the perspective of residue chemistry, the R→K→L substitution is accompanied by a decreasing positive charge density (R and K are positively charged, whereas L is hydrophobic); the guanidino group unique to R can form a multi-directional hydrogen-bond network through charge delocalization and, owing to its high pKa, maintains a positive charge more stably. This gradient is consistent with the activity gradient (DRE > DKE ≈ DLE), suggesting that this site may influence the local structural environment through side-chain charge and hydrogen-bonding properties and thereby be associated with activity.

#### 3.4.4 Structural localization and functional association of the key sites

To determine the functional significance of the key windows that are highly correlated with activity, they were mapped onto the three-dimensional structure of the Fe protein (NifH) (PDB: 1N2C [36]). Because the Fe protein exists as two symmetric dimers (chains E/F and G/H) in this crystal structure, chain F was selected as a representative NifH monomer for the structural mapping analysis. The three windows most strongly correlated with activity (977, 738 and 742) and the window with the strongest negative correlation (1026) correspond to residues 190–192, 53–55, 57–59 and 226–228 of NifH, respectively. Since the 12 involved positions have a 0% gap proportion across all 321 training sequences, their alignment columns directly and identically map to this same set of residue coordinates in the reference structure (PDB: 1N2C). It should be noted that this analysis addresses the spatial positions of these sites in the 3D structure, rather than the specific amino acid identities in individual sequences. For ease of discussion, NifH is divided here into four structural elements based on the definitions of Schlessman et al. [43]: the P-loop (Walker A, residues 9–16), Switch I (residues 38–43), Switch II (residues 125–135) and the 60’s loop (residues 60–70). All the numbering above follows the Azotobacter vinelandii Fe protein (Av2) sequence, consistent with chain F of PDB 1N2C. The minimum heavy-atom distances from the residues contained in each key window to the above structural elements and key ligands were calculated (**Figure 7**), and the results show that the four highly correlated windows converge spatially on two classes of functional interface of the Fe protein.

**Figure 7.**
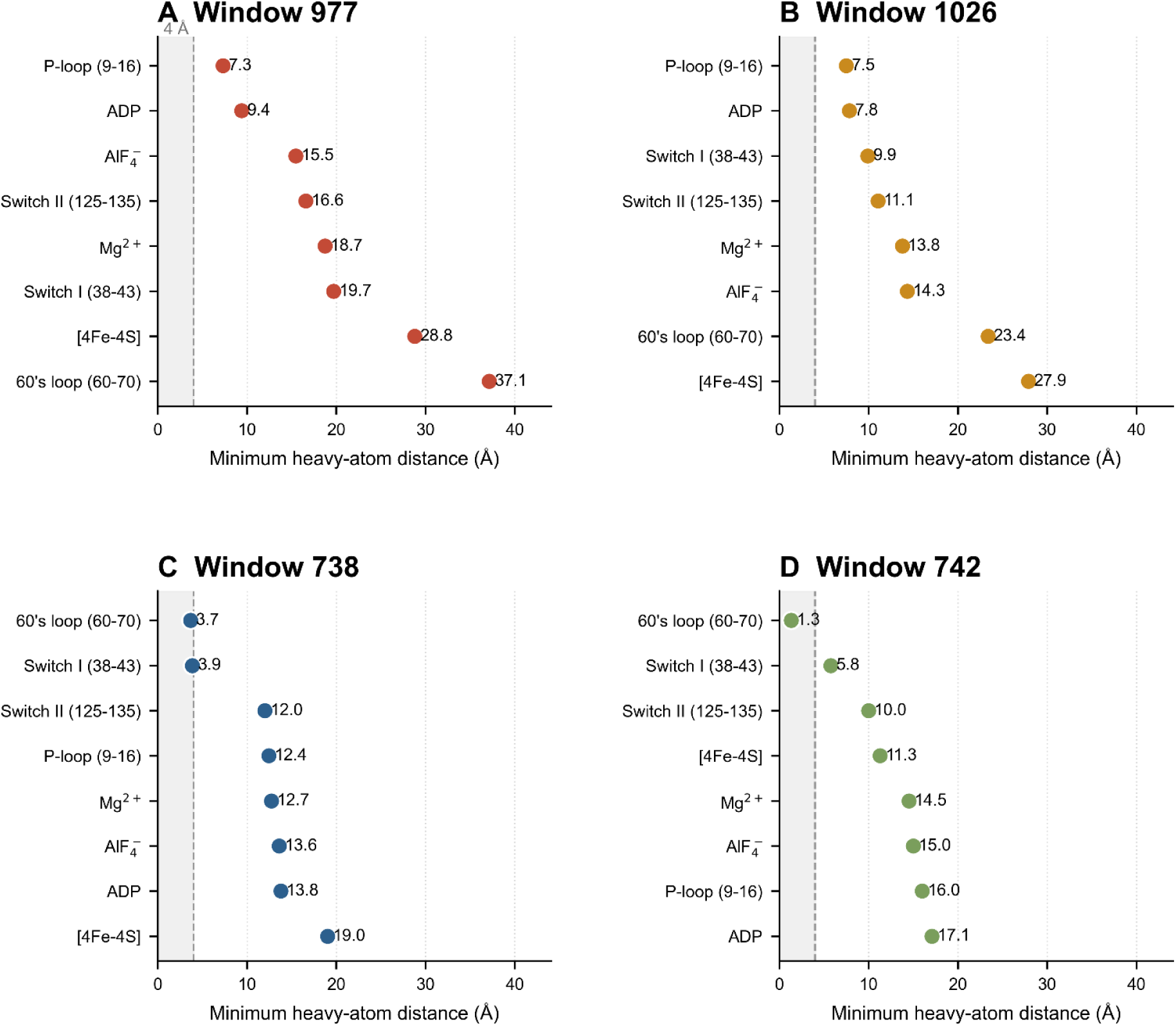
Spatial localization of the key sites in the NifH structure. (A–D) Minimum heavy-atom distances between the residues of each key window and the structural elements and key ligands of NifH (PDB: 1N2C, chain F). (A) Window 977 (residues 190–192); (B) window 1026 (residues 226–228); (C) window 738 (residues 53–55); (D) window 742 (residues 57–59). The *x*-axis gives the minimum heavy-atom distance (Å) between the window residues and each feature; features are ordered by increasing distance along the *y*-axis. The shaded region marks distances below 4 Å, used as a conservative upper bound for direct hydrogen-bonding or salt-bridge contact; the dashed line marks this 4 Å threshold. Minimum heavy-atom distances were computed from the 1N2C crystal structure, excluding hydrogen atoms.

First, the minimum heavy-atom distance of window 977 is 7.32 Å to the P-loop and 9.40 Å to ADP, but 28.79 Å to the [4Fe-4S] cluster; window 1026 shows a similar pattern, being 7.49 Å from the P-loop, 7.83 Å from ADP and 27.91 Å from the [4Fe-4S] cluster. Taking the 4 Å distance criterion for salt-bridge formation [44] as a conservative upper bound for direct interaction, all these distances exceed that range, suggesting that neither window has a direct electrostatic or hydrogen-bonding interaction with the P-loop or with bound ADP and that both are more likely related to it indirectly through intermediate residues or at the conformational level. The residues flanking window 977 are almost completely conserved in the training sequences (D at the first position in 99.1%, E at the third position in 100%), whereas its central position is polymorphic (R 61%, K 33%, L 4%). Because the flanks are invariant while the central position varies, and because this variation tracks activity (**Figure 6C**), we infer that the central side chain is functionally relevant. Given its >4 Å separation from both the P-loop and ADP, this influence is unlikely to be direct, and may instead propagate through the hydrogen-bonding network connecting this position to the nucleotide-binding site, thereby affecting the conformational change of NifH upon nucleotide binding, a mechanism consistent with the nucleotide-dependent conformational coupling observed for Switch-region residues in Fe protein [25]. Consistent with a role for hydrogen-bonding geometry rather than net charge, Arg and Lys carry the same charge yet differ in activity, whereas Lys and uncharged Leu do not (**Figure 6C**). It should be noted that the nucleotide bound in 1N2C is the ADP·AlF_4_^-^ transition-state analogue [36].

Second, the minimum distance of window 738 is 3.67 Å to the 60’s loop and 3.88 Å to Switch I, and window 742 is only 1.33 Å from the 60’s loop, immediately adjacent to its initial residue 60. The two windows are close to two structurally distinct regulatory elements—Switch I (a canonical nucleotide-dependent switch) and the 60’s loop (a protein-docking interface)—and may participate in the transmission of conformational changes and interface assembly. The 60’s loop is a hypervariable region of NifH located on the MoFe protein-binding surface, whose residues participate in stabilizing the Fe protein–MoFe protein complex [38, 43], suggesting that 738 and 742 may participate in the assembly of the electron-transfer complex by affecting conformational adjustments at this interface. The above distribution indicates that the key signals identified by the model can be structurally related to two processes, namely nucleotide binding (977, 1026) and conformational transmission/protein docking (738, 742).

It should be pointed out that this analysis is based on the static spatial relationships in a single representative structure. Although the key sites have no insertions or deletions in any of the training sequences, flexible regions such as the 60’s loop differ naturally in conformation among species [43], and a single structure can hardly represent the conformational diversity within the sequence space; the minimum heavy-atom distance also reflects only the spatial proximity in a static crystal structure and does not constitute direct evidence of dynamic coupling, so its molecular mechanism remains to be verified by site-directed mutagenesis and molecular dynamics simulations. Nitrogenase activity is also affected by factors such as environmental signals, cofactor state and post-transcriptional regulation, and analyses based only on sequence and static structure can hardly reflect its dynamic regulation.

#### 3.4.5 Conservation comparison of the key windows

This section further examines the conservation of these windows in the sequence. The conservation (the mean frequency of the most common residue at the three sites) and the mean description intensity were calculated for the 894 strict windows, and the two were significantly negatively correlated (Spearman ρ = −0.295, ***P*** = 2.1 × 10^−19^; **Figure 8**): the mean description intensity of highly conserved windows (conservation ≥ 0.9, *n* = 300) was 22.56, significantly lower than that of low-conservation windows (conservation < 0.7, *n* = 254; 27.80; Mann–Whitney ***P*** = 6.9 × 10^−18^). Correspondingly, among all 84 highly correlated windows (|*r*| > 0.3), only 7 windows had a conservation ≥ 0.9 (8.3% of *n* = 84), far below the proportion of 33.6% (*n* = 894) of windows with conservation ≥ 0.9 in the full set, which is consistent with the observation that the regions harboring absolutely conserved motifs of nitrogenase (such as the P-loop and the [4Fe-4S] cluster ligands of NifH, and the cofactor ligands of NifD and NifK [24, 40]) correlate poorly with activity. This negative result is not a defect of the model: absolutely conserved sites do not themselves carry information that distinguishes activity differences among strains, and the model therefore allocates its limited information capacity to variable sites that are associated with activity. This also explains the spatial distribution of the highly correlated windows—they are mostly located adjacent to functional elements rather than inside them: the residues at these positions are themselves variable and therefore discriminative, and can at the same time influence the conformation and function of conserved elements through spatial proximity, so that they are informative at both the sequence and the structural level.

**Figure 8.**
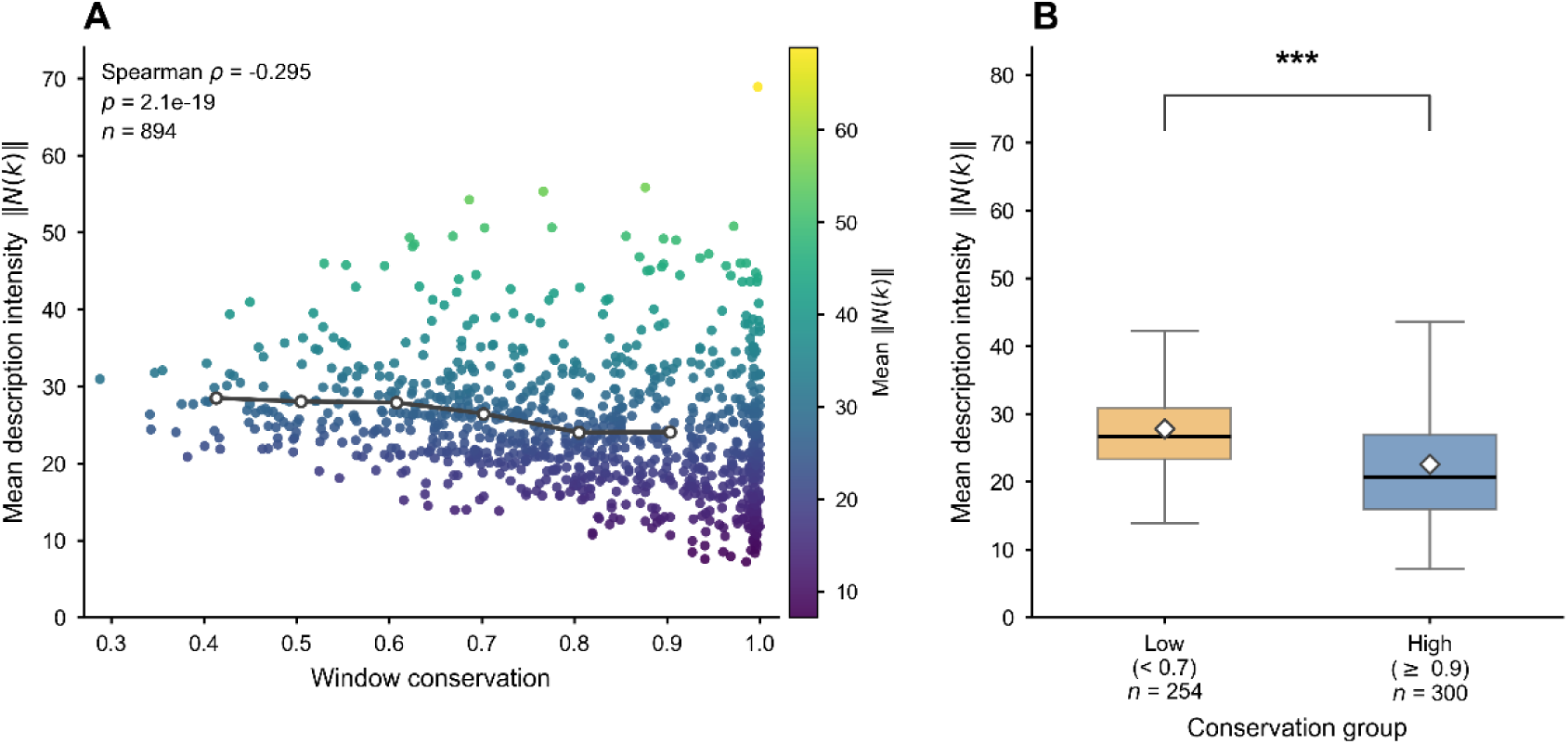
Conservation control for window-level description intensity. (A) Mean description intensity ‖N(k)‖ against window conservation for the 894 strict windows; the black line shows the moving average, and the color scale encodes mean intensity. Spearman *ρ* = −0.295, *p* = 2.1 × 10^-19^. (B) Comparison of mean description intensity between low-conservation (< 0.7, *n* = 254) and high-conservation (≥ 0.9, *n* = 300) window groups (Mann–Whitney U test, ***: *p* < 0.001). Absolute conservation of functional motifs carries no strain-discriminating information, and the model allocates its capacity to variable, activity-associated positions.

## 4. Conclusions

Following the Carmna work [22], this study proposes CarmnaDD, a purely sequence-driven framework based on the DD model for the classification and regression of nitrogenase activity. The framework encodes sequences into two sets of learnable parameters, the composition weight map (CWM) and the position weight function (PWF), requires no external priors such as pretrained protein language models, and learns representations end-to-end from the raw amino acid sequence alone. On the 402 nitrogenase sequences, CarmnaDD achieved *R*^2^ = 0.6035 in the regression task and AUC = 0.9377 and F1 = 0.8642 in the classification task, further improving the prediction performance on the benchmark established by Carmna and outperforming six classical machine learning baselines.

Beyond prediction performance, the internal representations of CarmnaDD show a systematic correspondence with known biological features of nitrogenase: the high-weight fragments captured by the CWM are significantly enriched in polar uncharged residues, consistent with the conclusion of Carmna based on SHAP analysis, reflecting common ground between the two modeling routes at the level of compositional preference; in addition, an enrichment trend of positively charged residues (K, R and H) was observed, which is in line with the structural biology knowledge that these residues participate through electrostatic interactions in cofactor assembly, subunit recognition and regulation of the catalytic microenvironment in nitrogenase. At the same time, the highly correlated sequence windows are structurally distributed around variable sites adjacent to two classes of functional interfaces, nucleotide binding and conformational transmission/protein docking, rather than inside absolutely conserved catalytic motifs, suggesting that the sequence information that distinguishes the activitys of strains resides mainly in regions that are functionally constrained but retain room for variation.

These results show that CarmnaDD achieves further performance improvement over Carmna with a lighter input, and that its internal representations show multiple correspondences with the known structural and functional features of nitrogenase, thus providing a computational framework that combines both performance and interpretability for nitrogenase activity prediction and functional site discovery, and an analysis strategy that can be extended to the study of nitrogen-fixing microorganisms for which expression data are scarce. In the future, combining site-directed mutagenesis with molecular dynamics simulations may further elucidate the molecular mechanisms of the key sites highlighted in this study.

## Data availability

The source code and datasets used in this study are available on GitHub (https://github.com/mbglab/DualDescriptor/tree/main/Apps/CarmnaDD).

